# VDAC2 stabilizes a membrane-inserted, primed intermediate of BAX activation

**DOI:** 10.64898/2026.07.13.738215

**Authors:** Varun Ravishankar, Laidy M. Alvero-González, Agathe Hubert, Zaiwei Zhang, Chloé Markarian, Valérie Prima, Sirine Chergui, Roberto Melero, Florian Stengel, Stéphen Manon, Jean-Pierre Duneau, Maria Queralt-Martin, Lucie Bergdoll

## Abstract

BAX, a major effector of mitochondrial apoptosis, is activated through a series of conformational transitions that lead to mitochondrial outer membrane permeabilization. Genetic studies have established VDAC2 as an essential regulator of BAX-mediated apoptosis, yet the molecular basis of this regulation remains unresolved. The absence of direct structural and biochemical characterization of VDAC2–BAX interactions has prevented mechanistic understanding of how VDAC2 influences BAX activation. Here, using complementary biochemical, biophysical and structural approaches, we reconstituted and characterized a stable VDAC2–BAX complex that was not observed with VDAC1, highlighting an isoform-specific role in BAX regulation. We show that VDAC2 captures and stabilizes a primed BAX conformation displaying the hallmarks of activation, including membrane insertion, BH3 exposure, and increased accessibility of the N-terminal activation region. By integrating AlphaFold3 predictions with molecular dynamics simulations, biochemical, biophysical and structural constraints, we derive an experimentally-supported structural model in which BAX is anchored through its α9 helix while its soluble domain partially extends over the VDAC2 pore. Together, our findings support a model in which VDAC2 facilitates BAX membrane insertion and stabilizes a membrane-inserted, activation-competent BAX intermediate. Rather than serving as a structural component of apoptotic pores, VDAC2 acts as a regulatory checkpoint in BAX activation. These results provide a molecular explanation for the emerging role of VDAC2 in mitochondrial apoptosis and establish structural basis for a previously inaccessible intermediate in the BAX activation pathway.

## Introduction

Apoptosis is a tightly regulated form of programmed cell death essential for development, tissue homeostasis, and the elimination of damaged or harmful cells. In the intrinsic pathway, mitochondrial outer membrane (MOM) permeabilization commits cells to death through the release of apoptogenic factors such as cytochrome c^1^. This process is controlled by the BCL-2 family of proteins, among which BAX and BAK act as the core pore-forming effectors^2,3^. While BAK is constitutively anchored at the MOM, BAX is predominantly cytosolic in healthy cells and must translocate to mitochondria upon activation^4,5^. BAX activation involves a series of conformational transitions, including exposure of the N-terminal activation epitope recognized by the conformation-specific antibody 6A7^6,7^, membrane insertion of the C-terminal helix α9 and exposure of the BH3 domain, ultimately leading to oligomerization and pore formation^8^. Structural studies have provided detailed views of soluble BAX, detergent-induced dimers complexed with BH3 peptides, and apoptotic pores^9–12^. Yet, the transient intermediates linking these states and the molecular mechanisms underlying BAX recruitment, membrane engagement, and oligomerization remain incompletely understood.

Among the mitochondrial factors implicated in these early events, VDAC2, originally known as an ion channel involved in ion and metabolite homeostasis, has emerged as an important contributor of BAX-dependent apoptosis^13–17^. Genetic ablation of VDAC2 profoundly modifies BAX-mediated cell death^13^, establishing VDAC2 as an essential component of the BAX activation process. VDAC2 also plays a critical role in the regulation of BAK, further highlighting its importance in mitochondrial apoptosis^14–21^. Despite this extensive genetic and cellular evidence, the molecular basis of VDAC2 function in apoptosis remains poorly understood. Previous studies have proposed that VDAC2 may participate in multiple stages of the BAX activation pathway, including mitochondrial recruitment, membrane insertion, and retrotranslocation ^14,22^. However, the molecular nature of the VDAC2-bound BAX species remains unknown. In the absence of direct biochemical and structural characterization, it is unclear whether VDAC2 engages BAX through transient encounters or stable complex formation, or how this interaction influences the conformational transitions that drive BAX activation and apoptotic pore formation. While mutagenesis and crosslinking studies have identified regions of VDAC2 involved in BAK binding and suggested partially overlapping interaction surfaces for BAX and BAK^13,17,19^, no experimentally validated structural model of a VDAC2–BAX complex is currently available. Characterizing the structure and functional properties of this complex is therefore essential for understanding how VDAC2 regulates the progression of BAX from a soluble monomer to a membrane-inserted pore-forming assembly.

Here, we sought to define the molecular basis of VDAC2-dependent BAX activation and to characterize a membrane-associated intermediate that precedes BAX oligomerization. Using complementary biochemical, biophysical, structural and molecular dynamics (MD) simulations approaches, we show that VDAC2 captures and stabilizes a primed BAX conformation displaying the three hallmarks of activation: α9 membrane insertion, BH3 exposure, and exposure of the N-terminal 6A7 epitope. Although this species displays the hallmarks of BAX activation, it remains pre-oligomeric and therefore represents a previously unidentified intermediate in the BAX activation pathway rather than the final pore-forming state. Together, our findings provide a mechanistic basis for previously proposed roles of VDAC2 in BAX recruitment, membrane insertion and retrotranslocation.

## Results

### BAX forms a stable complex with VDAC2

VDAC2 has been implicated in BAX activation, yet direct interaction between the two proteins has never been demonstrated at the molecular level. To determine whether VDAC2 can directly engage BAX and form a stable complex, we established a strategy to produce and purify VDAC2–BAX assemblies. Because BAX conformation is highly sensitive to its membrane environment—some detergents induce BAX activation, dimerization and exposure of the 6A7 epitope^23^, whereas cardiolipin-rich liposomes promote membrane insertion even in the absence of activators such as tBID^24^—we used a detergent-free nanodisc system composed of simple egg phosphatidylcholine lipids.

His-tagged VDAC2 was first reconstituted into lipid nanodiscs and included during the cell-free synthesis of untagged BAX (Fig. 1A). Following affinity purification through the VDAC2 His-tag, BAX efficiently co-eluted with VDAC2-containing nanodiscs (Fig. 1B). In contrast, BAX failed to co-elute with empty nanodiscs (Fig. 1B), as previously observed^25^. These results demonstrate that VDAC2 promotes stable association of BAX with membrane nanodiscs.

**Fig 1.**
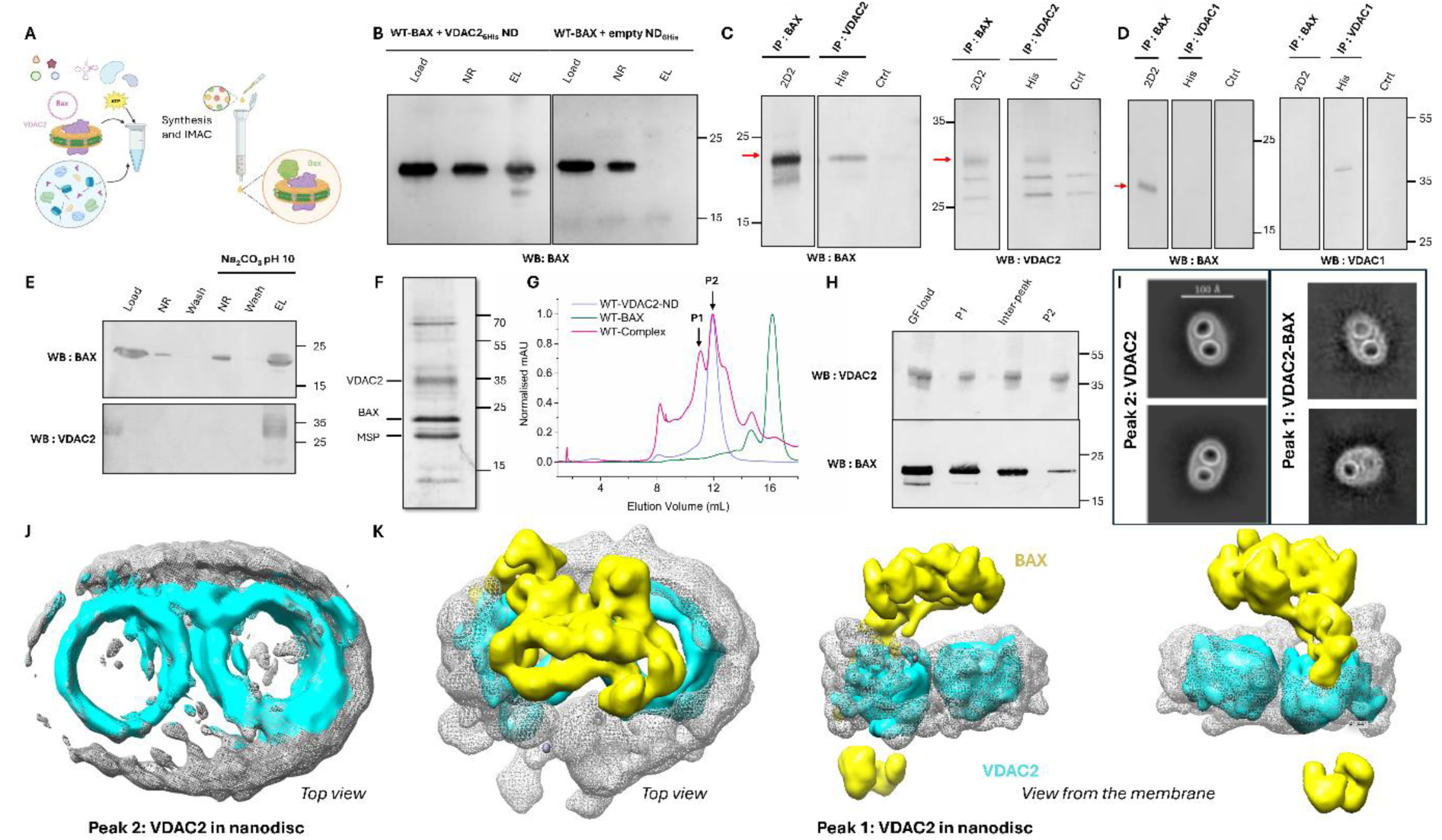
VDAC2 directly forms a stable membrane-associated complex with BAX **A.** Schematic of the experimental strategy. Full-length BAX was synthesized by cell-free expression in the presence of preformed His-tagged VDAC2-containing nanodiscs and purified by Ni-NTA affinity chromatography through the VDAC2 His tag. **B.** Western blot analysis of BAX following cell-free synthesis and affinity purification with 6His- VDAC2-containing nanodiscs or His-tagged empty nanodiscs. BAX co-elutes with VDAC2-containing nanodiscs but not with empty nanodiscs. **C.** Co-immunoprecipitation of WT BAX and VDAC2. **D.** WT BAX and VDAC1, following cell-free synthesis in the presence of VDAC2- or VDAC1-containing nanodiscs. Intact nanodiscs were immunoprecipitated using either a BAX-specific antibody (2D2) or an anti-His antibody recognizing VDAC2 and captured on Protein G Sepharose. After binding, the nanodiscs were solubilized with 0.5% NP-40 to disrupt the lipid bilayer while preserving direct protein–protein interactions. Resin without antibody served as a negative control. **E.** Sodium carbonate extraction (pH 10) of the purified VDAC2–BAX complex. Following cell-free synthesis in the presence of VDAC2-containing nanodiscs, complexes were immobilized on Ni-NTA resin through the VDAC2 His tag. After washing away unbound BAX, a sodium carbonate wash (pH 10) was used to detach peripherally associated BAX. The remaining membrane-associated complex was subsequently eluted with imidazole (EL). A fraction of BAX was released by sodium carbonate, whereas the majority remained associated with VDAC2-containing nanodiscs. **F.** SDS–PAGE analysis of the Ni-NTA-purified VDAC2–BAX complex showing VDAC2, BAX and the membrane scaffold protein MSP1D1. **G.** Size-exclusion chromatography profiles of VDAC2-containing nanodiscs (blue), soluble BAX (green) and the purified VDAC2–BAX complex (magenta). **H.** Western blot analysis of fractions collected from the size-exclusion chromatography of the VDAC2– BAX complex. **I.** Representative cryo-EM 2D class averages obtained from the final particle sets corresponding to SEC peak 2 (left) and SEC peak 1 (right). **J.** Ab initio cryo-EM reconstruction of a VDAC2 dimer reconstituted in an MSP1D1 nanodisc, obtained from particles corresponding to SEC peak 2. **K.** Ab initio cryo-EM reconstruction of a VDAC2 dimer reconstituted in an MSP1D1 nanodisc in complex with BAX, obtained from particles corresponding to SEC peak 1. VDAC2 is shown in blue, BAX in yellow, and the nanodisc density in grey.

We next asked whether BAX and VDAC2 form a genuine protein complex or are merely incorporated into the same nanodisc. To distinguish between these possibilities, the purified VDAC2–BAX complex in nanodiscs was first subjected to immunoprecipitation using antibodies against BAX (2D2) or the VDAC2 His tag. After capture on the Protein G-sepharose resin, the nanodiscs were solubilized with 0.5% NP-40 to disrupt the lipid bilayer and remove any association mediated solely by co-partitioning within the same nanodisc. Under these conditions, BAX and VDAC2 reciprocally co- immunoprecipitated (Fig. 1C), demonstrating that their association persists after disruption of the nanodisc membrane. Thus, VDAC2 and BAX form a stable protein complex rather than simply coexisting within the same nanodisc. Under identical conditions, BAX failed to co-immunoprecipitate with VDAC1, indicating that stable complex formation is specific to VDAC2 (Fig. 1D).

To assess the membrane state of BAX associated with the VDAC2 nanodisc, we performed sodium carbonate extraction at pH 10, which removes proteins peripherally associated with membranes while retaining membrane-inserted species. Following carbonate treatment, a fraction of BAX was detached from the nanodisc, whereas the majority of BAX remained associated with VDAC2-containing nanodiscs (Fig. 1E). These results suggest that the BAX associated to VDAC2-nanodiscs exists in two states, one resistant and one sensitive to carbonate extraction, consistent with the membrane-associated and membrane-inserted populations previously described in mitochondria ^14^.

Having established the existence of a stable VDAC2–BAX complex, we next sought to isolate and visualize this assembly. SDS-PAGE analysis of the purified sample confirmed the presence of VDAC2, BAX and MSP1D1, the scaffold protein used to form the nanodiscs (Fig. 1F). Size-exclusion chromatography revealed two major peaks: peak 1 containing both proteins, as confirmed by Western blot analysis, and peak 2, containing only VDAC2 nanodiscs (Fig. 1G,H). To visualize these assemblies, fractions corresponding to the SEC peaks were analyzed by cryo-EM. Two-dimensional class averages revealed distinct particle populations, with a subset of particles from peak 1 (VDAC2 and BAX in nanodisc) displaying partial occlusion of a VDAC pore (Fig. 1I). Three-dimensional ab initio reconstructions were generated for both VDAC2 nanodiscs alone and VDAC2–BAX nanodiscs (Fig. 1J,K). Ab initio three-dimensional reconstructions of peak 2 showed the characteristic morphology of a VDAC2 dimer embedded in a lipid nanodisc (Fig. 1J), consistent with previous observations that VDAC2 has a strong tendency to dimerize in lipid bilayers^26,27^. In contrast, the reconstruction obtained from peak 1 contained a VDAC2 dimer in which one pore was partially obscured by additional density (Fig. 1K). Relative to the VDAC2-only reconstruction, two extra features were observed: a density positioned above one pore on the cytosolic side and a second density within the membrane. Although the resolution does not allow definitive assignment, these features are consistent with a membrane-inserted BAX molecule associated with VDAC2.Together, these observations demonstrate that VDAC2 and BAX form a stable membrane-associated complex that can be purified and structurally characterized, providing basis for subsequent functional and mechanistic analyses.

### Structural modeling and crosslinking identify α9-mediated engagement of BAX by VDAC2

Having established the existence of a stable VDAC2–BAX complex, we next sought to determine how BAX engages VDAC2 at the molecular level. Although mutagenesis and crosslinking studies have implicated the β10–β11 region of VDAC2 in BAK regulation^17,19^ and identified residues within the β6– β7 region as important for BAX interaction^13^, the molecular basis of BAX recognition by VDAC2 remains unknown, and no structural model of a VDAC2–BAX complex has been reported. We therefore combined structural modelling, crosslinking and mutagenesis to define the interaction interface and derive an experimentally constrained model of the complex.

To obtain an initial structural model of the VDAC2–BAX complex, we first employed AlphaFold-based structure prediction. As a validation of the approach, AlphaFold accurately reproduced the known inactive structure of BAX (Fig. 2A, SFig. S1A, RMSD = 2.46 Å +/- 0.26 Å with the NMR structure), providing confidence for subsequent modeling of the VDAC2–BAX complex. Initial AlphaFold2 predictions using VDAC2 and BAX consistently failed to identify the membrane region and to converge on a reproducible VDAC2–BAX complex. In contrast, AlphaFold3, which allows inclusion of fatty acids to mimic the hydrophobic environment of the membrane, generated convergent models across multiple independent predictions. In all cases, BAX engaged VDAC2 through its C-terminal α9 helix, which docked along β-strands β6–β7 of the VDAC2 barrel (Fig. 2B, SFig. S1B). Notably, the predicted interface encompasses residue A171 and strands β6–β7 of VDAC2, previously implicated in BAX regulation^13,17^. These observations support the predicted binding mode and suggest a direct membrane-embedded interaction mediated by the BAX α9 helix.

**Fig. 2:**
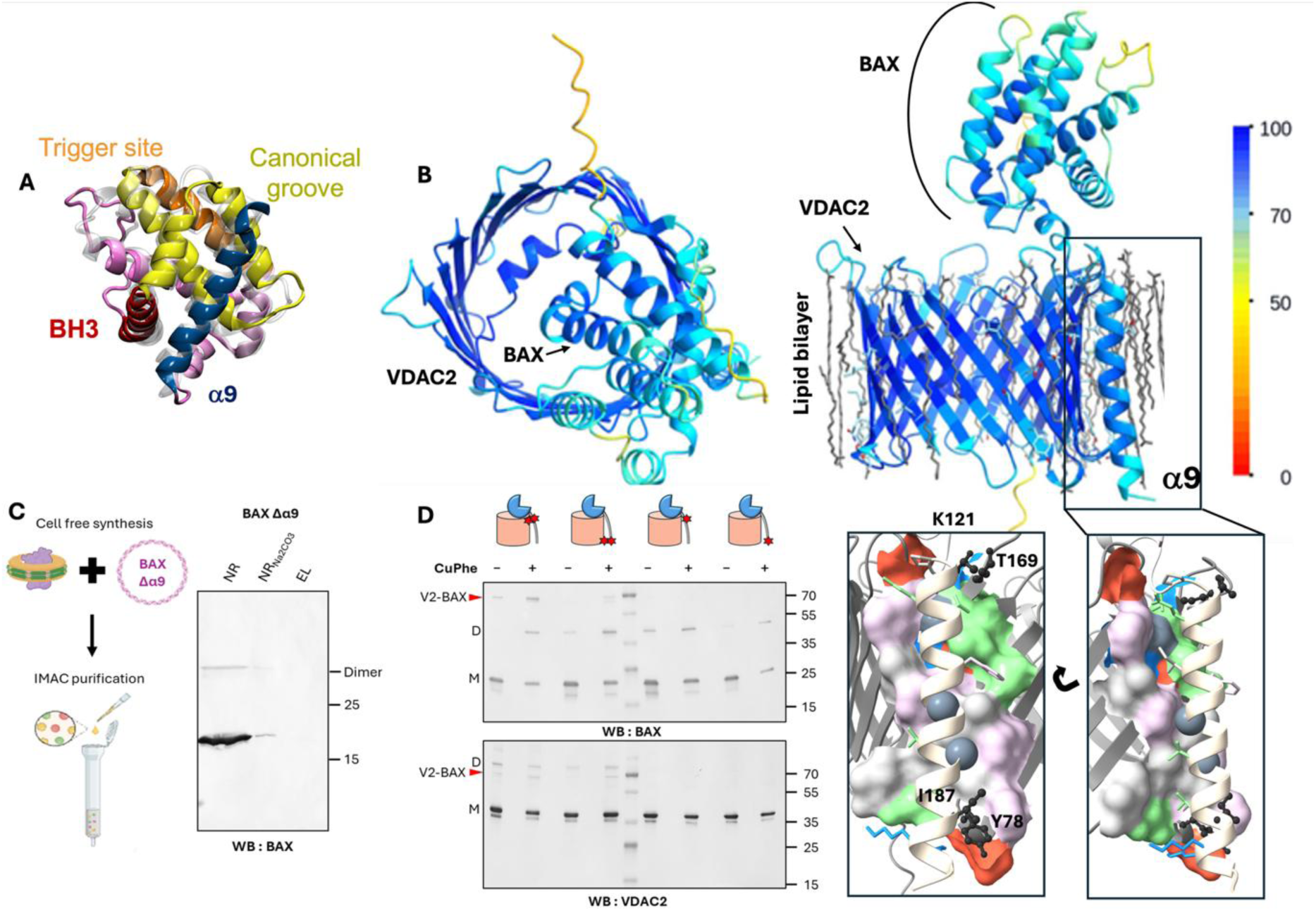
Structural modeling and crosslinking identify α9-mediated engagement of BAX by VDAC2. **A.** Structure of soluble inactive BAX (PDB: 1F16) highlighting the trigger site (orange), canonical groove (yellow), BH3 domain (red), and C-terminal α9 helix (blue). The AlphaFold3 prediction, shown in transparent grey, closely reproduces the experimental structure (RMSD = 2.46 Å +/- 0.26 Å; SFig. S1A). **B.** Representative AlphaFold3 model of the VDAC2–BAX complex colored according to pLDDT confidence values (scale bar on the right). 15 models from 3 independent predictions converged on the same α9-mediated interaction interface (SFig. S1B). **C.** BAX western-Blot after Ni-NTA affinity purification of VDAC2 nanodiscs following cell-free expression of BAX-Äá9 shows that BAX-Äá9 does not associate with VDAC2. **D.** CuPhe-mediated cysteine cross-linking of VDAC2 and BAX single-cysteine mutants validates the predicted α9-mediated interface. Right insets show close-up views of the predicted interaction surface. VDAC2 is displayed as a surface representation colored according to residue properties: acidic (red), basic (blue), uncharged polar (green), aliphatic (white), and aromatic (light pink). BAX helix α9 is shown in cartoon representation, with selected side chains displayed as sticks to illustrate their complementarity with the VDAC2 surface. Helix α9 contains a conserved GXXXA motif; the Cα and Cβ atoms of G179 and A183 are shown as grey van der Waals spheres. Residues mutated to cysteines for cross-linking validation (VDAC2 K121C/BAX T169C and VDAC2 Y78C/BAX I187C) are shown as black sticks. Left: Experimental validation of the α9 orientation by cysteine cross-linking. Western blots of complexes containing, from left to right, VDAC2 K121C/BAX T169C, VDAC2 Y78C/BAX I187C, VDAC2 ΔCys/BAX T169C, and VDAC2 ΔCys/BAX I187C. M and D indicate the monomeric and dimeric forms of each protein, respectively. The cross-linked VDAC2–BAX complex is indicated by a red arrow.

To assess the reliability of the predicted complex, we examined the confidence scores (pLDDT) and convergence across independent AlphaFold3 predictions. High confidence scores and excellent agreement between predictions were consistently observed for the VDAC2 β-barrel and the BAX α9 helix, supporting a robust membrane-embedded interaction interface (Fig. 2B, SFig. S1B). In contrast, the position of the soluble BAX domain relative to VDAC2 varied between models, despite the BAX globular domain itself remaining structurally conserved. These observations suggest that while the α9- mediated interaction is well defined, the soluble domain retains conformational flexibility and cannot be positioned unambiguously from AlphaFold3 predictions alone. We therefore considered the AF3 model as an initial structural hypothesis that required experimental validation and refinement.

To experimentally test the predicted interface in the membrane region, we introduced cysteine substitutions at residue pairs predicted to lie in close proximity (close up in Fig. 2D) and performed CuPhe-mediated crosslinking. Efficient cysteine crosslinking was observed for both VDAC2 K121C– BAX T169C and VDAC2 Y78C–BAX I187C pairs (Fig. 2D), consistent with the distances predicted by the model. To further assess the importance of α9, we examined a BAX construct lacking the last 22 amino acids (BAX-Δα9). In contrast to full-length BAX, BAX-Δα9 failed to form a stable complex with VDAC2 (Fig. 2C), demonstrating that α9 is required for stable engagement of BAX by VDAC2.

Together, structural modelling, crosslinking and mutagenesis identify a robust α9-mediated interaction interface that provides the foundation for subsequent experimental refinement of the VDAC2–BAX structural model. Analogous modelling of VDAC1 did not yield a convergent and reliable interaction interface (SFig. S1C), suggesting that stable complex formation is a specific feature of VDAC2, consistent with the biochemical data (Fig. 1D).

### The VDAC2–BAX complex is stable in membranes and partially occludes the VDAC pore

Having established a preliminary structural model for the VDAC2–BAX complex, we next asked whether this interaction could be maintained in a membrane environment and whether it might affect VDAC2 channel function. To address these questions, we first performed MD simulations of the complex embedded in a lipid bilayer.

Throughout three independent simulations of 10 μs, the overall architecture of the complex remained stable (Movies 1-3). The α9 helix remained associated with the same region of the VDAC2 barrel, supporting the robustness of the predicted interaction interface. In contrast, the soluble domain of BAX sampled multiple orientations relative to VDAC2 (representative conformations are shown in Fig. 3A).

**Fig. 3:**
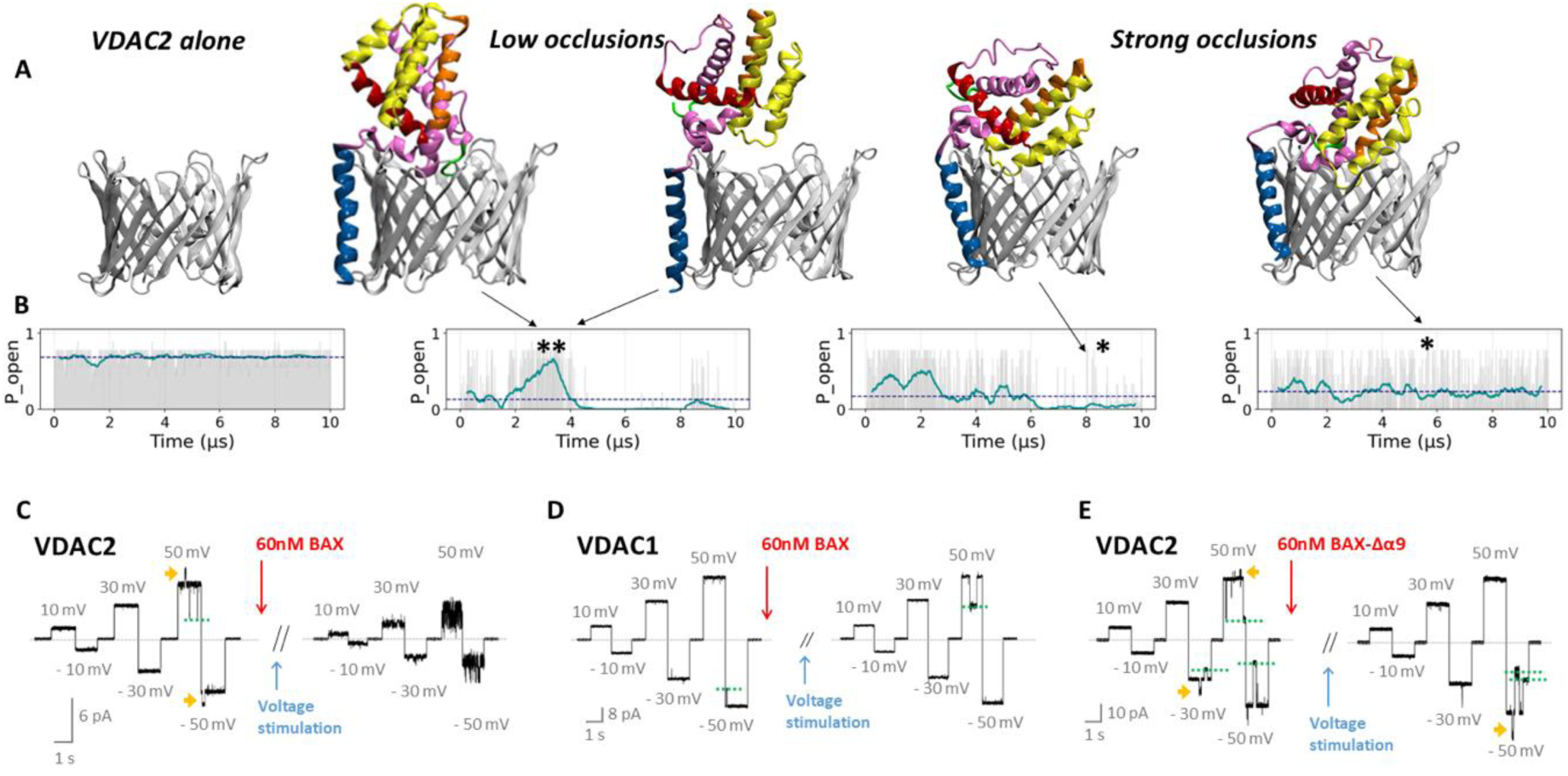
Molecular dynamics simulations and electrophysiology reveal stable α9-dependent pore occlusion by BAX. **A.** Representative snapshots from coarse-grained MD simulations (Movies 1-3) illustrating different conformations of the VDAC2–BAX complex and the relative positioning of the soluble BAX domain with respect to the VDAC2 pore. VDAC2 is colored in grey, BAX in pink with the trigger site (orange), canonical groove (yellow), BH3 domain (red), and C-terminal α9 helix (blue). **B.** Estimated pore opening probability for the VDAC-BAX complex during three independent 10-µs MD simulations. For comparison, pore opening probability was recalculated from the trajectory shown in **A** after computational removal of BAX from the simulation (left). Grey bars are the conductance for each snapshot. The continuous green line is the floating average over 50 ns. Dotted line is the average of computed conductance of open pore probability over the whole simulation. **C. D. E.** are representative current recordings from a single VDAC channel before (left) and after (right) addition of 60 nM recombinant full-length monomeric BAX to the cis side of the membrane **C.** VDAC2 ; **D.** VDAC1 ; **E.** VDAC2 with BAX Δα9. Green dotted lines show VDAC-characteristing gating events^28^.Yellow arrows indicate the characteristic short-lived open-conductance substates of VDAC2^29^. Applied voltages are shown in grey and the grey dashed line indicates the zero-current level. Stable current reduction was observed only after application of a voltage stimulation in the form of a slow triangular voltage ramp (5 mHz, ±60 mV) for at least 30 min (SFig. S2A). Current traces were digitally filtered at 500 Hz using an 8-pole low pass Bessel filter. Experiments were performed with VDAC inserted in a soybean polar lipid extract (PLE) membrane with bathing solutions of 150 mM KCl buffered with 5 mM HEPES at pH 7.4.

To assess the functional consequences of BAX binding, we estimated pore conductance from conformations sampled throughout the MD trajectories. These calculations revealed a reduction in conductance relative to VDAC2 alone (Fig. 3B). Inspection of the corresponding structures showed that BAX partially occludes the VDAC pore and that different orientations of the soluble BAX domain generate distinct degrees of obstruction, resulting in multiple metastable conductance states (Fig. 3A,B).

To experimentally test this prediction, we reconstituted purified hVDAC2 into planar lipid bilayers and monitored single-channel activity before and after addition of monomeric BAX. Following insertion into the membrane, hVDAC2 displayed the characteristic conductance and voltage-dependent gating behavior previously reported for this isoform (Fig. 3C)^28,29^. Addition of monomeric BAX did not immediately alter channel activity. Instead, after several minutes of voltage stimulation (SFig. S2A), the channel transitioned to reduced-conductance states (Fig. 3C). Once established, these states persisted throughout the recording and the original open-state conductance could not be recovered within the experimental time window (>1 h), indicating the formation of a remarkably stable BAX–VDAC2 complex. Importantly, multiple long-lived reduced-conductance states were observed, consistent with the distinct pore occlusion states predicted by MD simulations (SFig. S2B). These states are distinct from the characteristic voltage-dependent gating of VDAC, as they were essentially irreversible over the course of the recordings and exhibited substantially greater current noise. Similar conductance reductions were observed under different ionic strengths and lipid compositions, indicating that the interaction is robust to changes in the environment. However, addition of monomeric BAX to hVDAC1 failed to induce comparable long-lived conductance reductions, indicating that pore-associated complex formation is specific to VDAC2 (Fig. 3D).

To determine whether α9 is required for this effect, we repeated these experiments using BAX-Δα9. In contrast to full-length BAX, the Δα9 mutant failed to induce stable channel blockades (Fig. 3E). Furthermore, the use of dimeric BAX, in which α9 is sequestered in the dimerization interface^30^, failed to interact with VDAC2 (SFig. S2C). Both experiments demonstrate that α9-mediated engagement is required for functional interaction with VDAC2.

Together, MD simulations and electrophysiological recordings support a model in which monomeric BAX forms a stable α9-dependent complex with VDAC2 that partially occludes the channel pore.

### VDAC2 stabilizes a primed BAX intermediate

The structural and functional analyses described above indicate that VDAC2 forms a stable complex with monomeric BAX. We next sought to determine the conformational state of BAX within this complex. BAX activation is characterized by three major structural hallmarks: insertion of the C- terminal α9 helix into the membrane, exposure of the BH3 domain, and structural changes at the N- terminus leading to exposure of the 6A7 epitope. Having established α9 insertion in the previous sections, we investigated whether the two remaining activation markers were also present in the VDAC2-bound state and could be used to refine our structural model.

Exposure of the BH3 domain is a critical step in BAX activation, as it enables the intermolecular interactions required for oligomerization. To probe BH3 accessibility, we performed NEM–PEG labeling experiments on the two native cysteines of BAX (C62 and C126) and on a cysteine introduced within the α9 helix (V177C). Compared with soluble BAX, labeling of C126, located in the α5–α6 loop, remained unchanged, indicating that this region is accessible in both conformational states. In contrast, V177C was largely inaccessible in the VDAC2-bound complex, consistent with the proposed model in which the α9 helix is inserted into the membrane. Strikingly, C62, located within the BH3 domain, was poorly accessible in soluble BAX but became readily labeled upon complex formation with VDAC2, indicating increased exposure of the BH3 region (Fig. 4A).

**Fig. 4:**
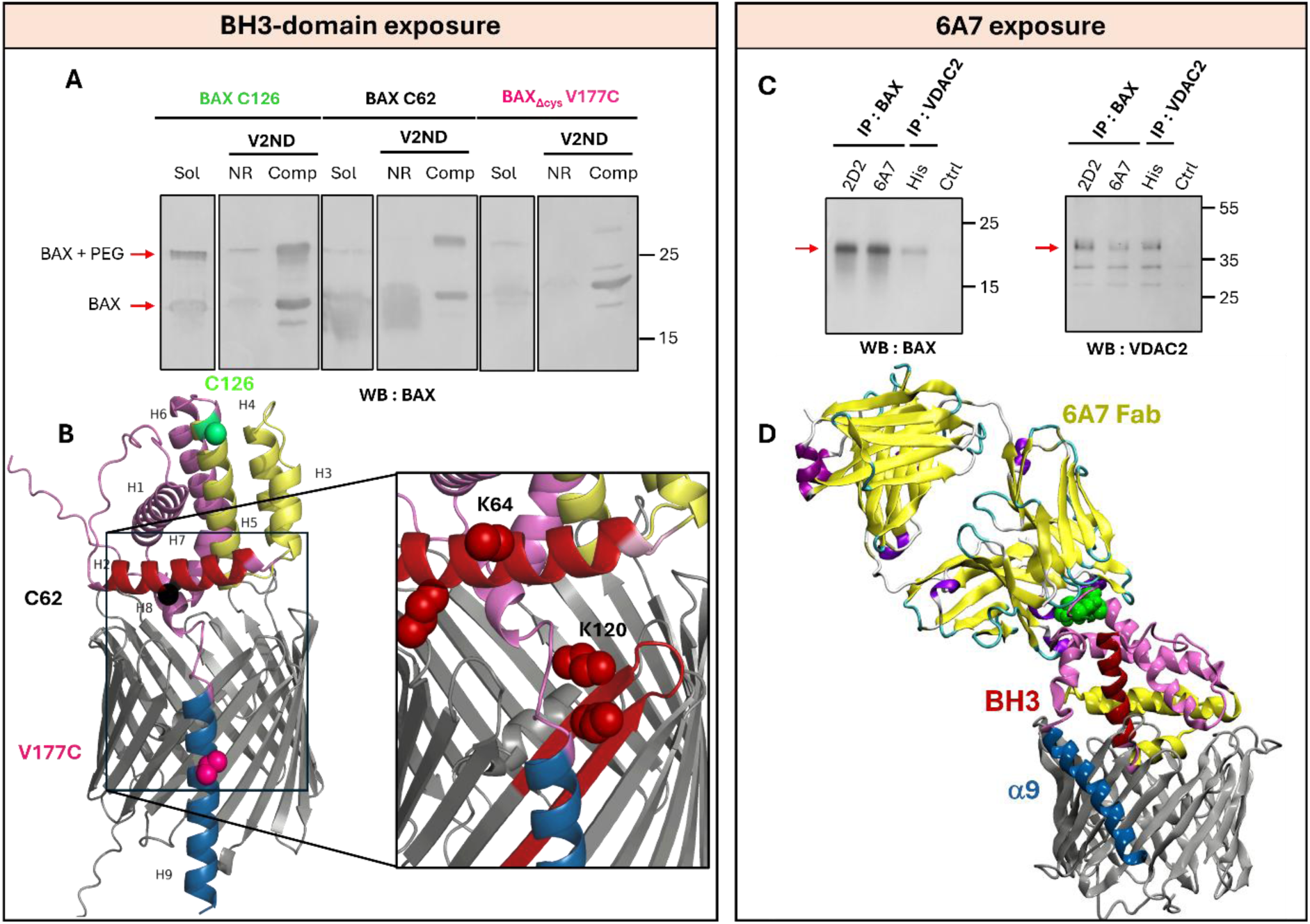
Integrative structural analysis reveals a membrane-inserted primed conformation of BAX bound to VDAC2. **A.** Accessibility of cysteine residues in soluble BAX and VDAC2-bound BAX probed by NEM-PEG labeling. Single-cysteine BAX variants (C122, green; C62, black; C177, magenta), shown on the structure below, were produced by cell-free expression in the presence of preformed WT VDAC2- containing nanodiscs, purified by affinity chromatography, and labeled with NEM-PEG (5 kDa). Labeling was detected by western blotting using the pan-BAX antibody 2D2. PEGylated species, corresponding to solvent-accessible cysteines, display reduced electrophoretic mobility. Sol: soluble BAX produced in the absence of VDAC2. NR: non-retained fraction after Ni-NTA purification of BAX produced in the presence of 6His-VDAC2 nanodiscs. Comp: BAX in complex with VDAC2 nanodiscs, eluted with imidazole. **B.** Model of the VDAC2-BAX complex showing the positions of the three cysteines used in the NEM-PEG assay. VDAC2 is shown in grey and BAX in pink, with the canonical groove (yellow), BH3 domain (red), and C-terminal α9 helix (blue). The Cα atoms of the engineered cysteines are shown as spheres. In the inset, intermolecular cross-links identified by DSS cross-linking coupled to mass spectrometry (XL-MS) are highlighted in red. XL-MS identified contacts between the β6–β7 region of VDAC2 (Lys120 and Lys121) and the BH3 domain of BAX (Lys58 and Lys64). Reactive lysines are shown as spheres. **C.** Co-immunoprecipitation of WT BAX and 6His-VDAC2 following cell-free expression in the presence of VDAC2-containing nanodiscs using the pan-BAX antibody 2D2, the activation-specific antibody 6A7, or an anti-His antibody recognizing VDAC2. Resin without antibody was used as a negative control. **D.** Docking of the 6A7 Fab onto representative conformations of the VDAC2–BAX complex identifies models compatible with antibody binding and provides an additional structural constraint for refinement of the integrative model (SFig. 4). VDAC2 is shown in grey and BAX in pink, with the 6A7 epitope as green spheres, the canonical groove (yellow), BH3 domain (red), and C-terminal α9 helix (blue).

**Figure 5:**
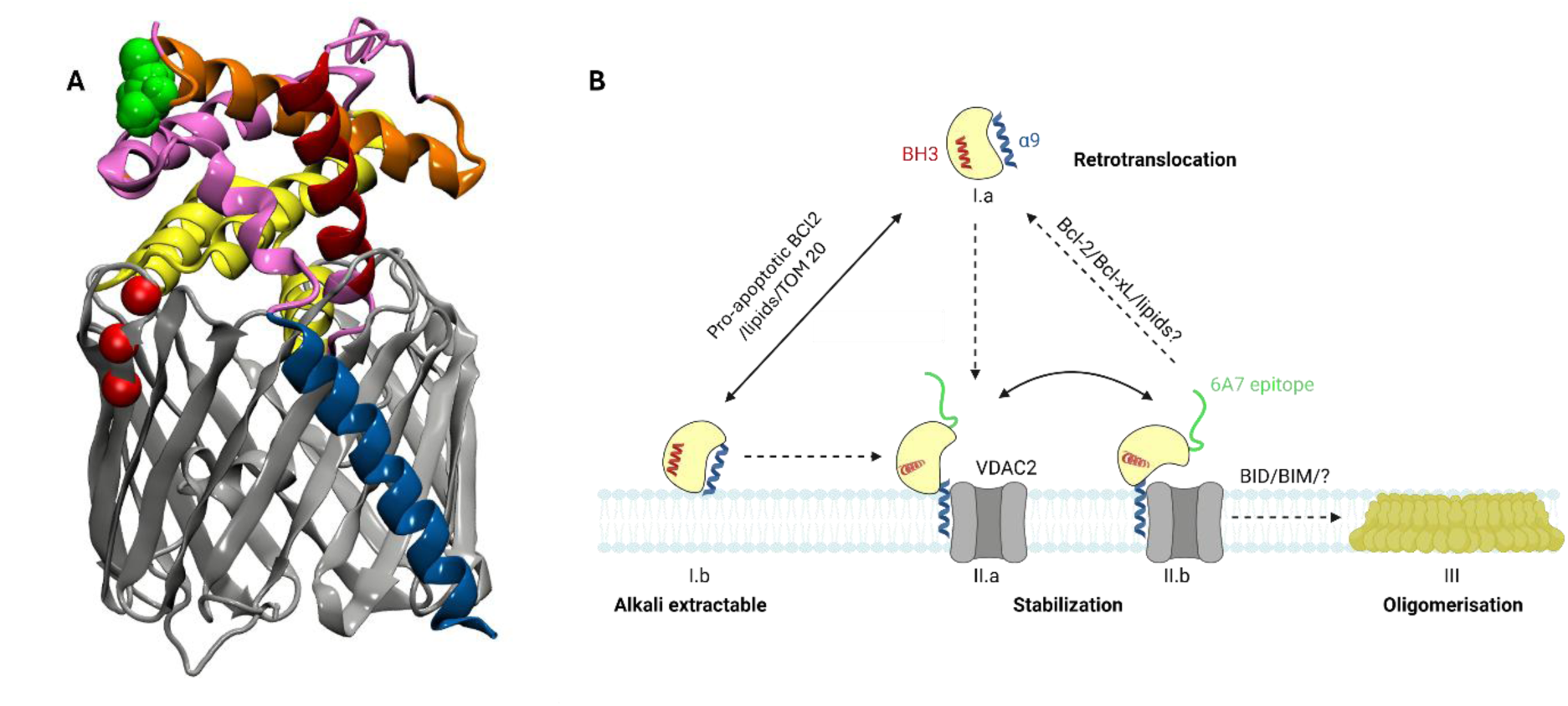
Structural model and functional role of the VDAC2–BAX complex. **A.** Integrative structural model of the VDAC2–BAX complex using MD simulations, cysteine crosslinking, XL-MS, cryo-EM, electrophysiology, mutagenesis, and antibody accessibility measurements. VDAC2 is shown in grey and BAX in pink, with the 6A7 epitope as green spheres, the canonical groove (yellow), BH3 domain (red), and C-terminal α9 helix (blue). VDAC2 residues previously implicated in VDAC2 regulation of BAX (A171)^17^ or BAK (T167 and D169)^19^ are highlighted as red spheres. **B.** Proposed role of VDAC2 during BAX activation. Following membrane engagement and α9 insertion, VDAC2 captures and stabilizes a membrane-inserted, activation-competent BAX state characterized by BH3 exposure and accessibility of the 6A7 epitope while remaining pre-oligomeric. This intermediate may subsequently progress toward BAX oligomerization and apoptotic pore formation or undergo deactivation and retrotranslocation, possibly with a BCl2-family anti-apoptotic^47^.

To independently assess the organization of this region, we performed DSS crosslinking coupled to mass spectrometry. An intermolecular crosslink connected the β6–β7 region of VDAC2 to the BH3 domain (Fig. 4B, SFig. 3), compatible with the predicted complex architecture (Fig. 4D). Together, these results indicate that the BH3 region becomes accessible upon VDAC2 binding and provide experimental support for the proposed structural model.

We next examined accessibility of the N-terminal activation region using the conformation-specific antibody 6A7, which recognizes an epitope exposed upon BAX activation^23^. VDAC2-bound BAX was efficiently recognized and immunoprecipitated by 6A7, with an efficiency comparable to that of the conformation-independent antibody 2D2 (Fig. 4E). These results indicate that essentially all VDAC2- bound BAX adopts a 6A7-positive activated conformation.

Beyond serving as a biochemical marker of activation, 6A7 recognition provides an independent structural constraint on the organization of the VDAC2-bound complex. We therefore evaluated different representative conformations obtained during MD simulations for their compatibility with antibody binding^31^. While docking of the 6A7 Fab onto the “low occlusion” conformations (Fig. 3A) was unsuccessful, a restricted subset of the “strong occlusion” conformations sampled during the simulations allowed unobstructed access to the epitope without steric clashes with VDAC2 or the membrane (Fig. 4F and SFig. S4). Notably, these conformations converged on a common arrangement in which the soluble BAX domain occludes the VDAC2 pore, consistent with the electrophysiological measurements, while remaining tethered to the membrane through its α9 helix.

Collectively, the α9-mediated interface identified by AlphaFold3, MD simulations, cysteine crosslinking and mutagenesis, the conductance changes predicted computationally and observed experimentally, the BH3 accessibility revealed by NEM-PEG labeling and XL-MS, and the 6A7 accessibility constraint converge to define an experimentally validated structural model for the VDAC2– BAX complex. Our final model integrates multiple orthogonal experimental constraints and narrows the conformational landscape to a limited number of architectures compatible with all available data. In these models, VDAC2-bound BAX displays the hallmarks of activation while remaining associated with VDAC2 as a stable pre-oligomeric complex, supporting the existence of a primed intermediate poised for further activation.

## Discussion

### VDAC2 directly captures and stabilizes BAX at the mitochondrial membrane

How BAX is recruited to and activated at the mitochondrial outer membrane remains a central unresolved question in apoptosis research. Although genetic and cellular studies have firmly established VDAC2 as an important contributor of BAX-dependent apoptosis^13–16^, the molecular basis of this relationship has remained unclear. Here, we demonstrate that BAX inserts in the membrane and forms a stable complex with VDAC2 in both nanodiscs and planar lipid bilayers, providing the first direct biochemical evidence that VDAC2 can engage BAX independently of additional mitochondrial components. These findings provide a molecular explanation for the observation that loss of VDAC2 phenocopies loss of BAX^13^ and place earlier reports of VDAC2-dependent BAX mitochondrial localization and retrotranslocation^14^ into a structural context. While previous studies have established that BAX can reach the mitochondrial outer membrane through multiple pathways involving BH3-only proteins or membrane interactions^32–37^, our reconstituted system demonstrates that VDAC2 alone can directly insert and stabilize BAX in the membrane.

The interaction is remarkably stable: the complex persisted throughout the electrophysiology recordings lasting several hours, contrasting with the transient interactions reported between VDAC channels and proteins such as tubulin or α-synuclein^29,38–40^ and distinguishes VDAC2 from Tom22, which can facilitate BAX membrane insertion but does not remain associated thereafter^36^.

### Structural basis of BAX engagement by VDAC2

To understand the molecular basis of this VDAC2-bound BAX intermediate, we integrated AlphaFold3 predictions with MD simulations, cysteine crosslinking, mutagenesis, XL-MS, cryo-EM, electrophysiology and antibody accessibility measurements to derive an experimentally constrained model of the VDAC2–BAX complex. The initial AF3 predictions served as a starting model that was progressively refined using independent experimental constraints.

AF3 predictions, cysteine crosslinking, α9 deletion and the inability of dimeric BAX–in which α9 helix is sequestered in the dimerization interface–to bind VDAC2 all identify the C-terminal α9 helix as a central determinant of engagement. These findings extend the established role of α9 in mitochondrial targeting^16,41^ and membrane insertion by identifying it as a key determinant of productive interaction with VDAC2: the predicted interface positions a conserved GXXXA motif of α9 (Fig. 2 close up), a motif frequently involved in membrane protein interactions^42^, against a complementary surface of the VDAC2 β-barrel, providing a plausible structural explanation for the remarkable stability of the complex.

Although α9 provides the primary membrane anchor, multiple observations indicate that stable complex formation also involves the soluble BAX domain. Cryo-EM reveals additional density above the cytosolic side of the pore, XL-MS identifies contacts between the BH3 region of BAX and VDAC2, and previous work showed that preventing BH3 exposure also disrupts VDAC2 binding^14^, suggesting an important contribution of the globular domain. Notably, BAX complexed with VDAC2 exposes its BH3 domain in the absence of any known activator such as BIM or tBID. Interestingly, available data on BAK point to a similar architecture: although the α9 helix is required for membrane insertion, additional contacts involving the soluble domain and VDAC2 loop regions have also been implicated in stable complex formation^17,43^. Incorporation of these experimental constraints, together with the accessibility of the 6A7 epitope, substantially refines the initial AF3 predictions and consistently positions the globular domain of BAX above and in interaction with the loops of the VDAC2 pore.

The refined model is remarkably consistent with the literature on both BAX and BAK interactions with VDAC2. The predicted interface overlaps the β6–β10 region previously implicated in BAX-dependent apoptosis^13^, places the BAX-specific residue A171 identified by Yuan and colleagues ^17^ directly adjacent to the interaction surface, and localizes previously identified BAK-binding residues to the same region of VDAC2^19^. Together, these observations provide a structural explanation for why BAX and BAK engage overlapping surfaces on VDAC2, as observed previously^17^.

### VDAC2 stabilizes a primed BAX intermediate

Our biochemical, structural and functional data indicate that VDAC2-bound BAX displays the major hallmarks of activation, including membrane insertion of α9, exposure of the BH3 domain and increased accessibility of the N-terminal activation region. However, it remains stably associated with VDAC2 and shows no evidence of oligomerization. Thus, we refer to this state as a primed intermediate, a conformation that has acquired the hallmarks of BAX activation yet remains pre-oligomeric and has not assembled into the apoptotic pore. To our knowledge, this represents the first structural description for such an intermediate, in which BAX is anchored to VDAC2 through its α9 helix while its globular domain rests above the pore.

Because VDAC is absent from apoptotic pores^12,44,45^, progression toward oligomerization likely requires release of BAX from VDAC2. In this framework, VDAC2 acts as a stabilizing platform for a primed activation intermediate that precedes pore formation. Alternatively, since it has been demonstrated that BAX conformational changes are reversible, BAX associated with VDAC2 might provide a platform for its retrotranslocation as previously suggested^14^. VDAC2 might thus act as a mitochondrial checkpoint at which the cell fate is determined.

The interface identified here may also help explain the broader role of VDAC2 in regulating BCL-2 family proteins. Although BAX and BAK likely engage related surfaces on VDAC2 (^13,17,19^ and our model), the functional consequences of these interactions appear distinct. Whereas BAK is constitutively membrane-associated and is thought to be restrained by VDAC2 under non-apoptotic conditions^5,20^, our data indicate that VDAC2 captures BAX during its activation process and stabilizes a membrane-inserted, primed intermediate. VDAC2 may therefore regulate BAX and BAK at different stages of their activation pathways while using overlapping interaction surfaces, highlighting a specialized role in coordinating mitochondrial apoptosis beyond its canonical function as a metabolite channel.

Although our refined model places the globular domain partially over the VDAC2 pore and electrophysiological recordings reveal stable partial channel occlusion, this interaction is unlikely to significantly impair mitochondrial metabolite exchange in vivo. Mammalian cells contain several million VDAC molecules but only ∼10^5 BAX molecules^46^, implying that only a small fraction of channels could be occupied even if all BAX molecules were VDAC2-bound. We therefore favor a model in which the primary function of the VDAC2–BAX interaction is regulation of apoptosis rather than metabolite transport.

### Isoform-specific regulation of BAX by VDAC2

Under identical experimental conditions, VDAC1 failed to form a stable complex with BAX, did not support formation of the same activation-competent state, and did not produce the characteristic long-lived channel blockade observed with VDAC2. Notably, the complete absence of detectable interaction with VDAC1 contrasts with previously characterized VDAC interactors, including tubulin and α- synuclein, which display quantitative differences in binding among isoforms rather than a complete on/off behavior^29,38^. Together with previous genetic and cellular studies, these findings indicate that VDAC2 possesses a specialized capacity to capture and stabilize membrane-associated BAX that cannot be fulfilled by other VDAC isoforms^13,19,20^. Our model provides a molecular explanation for the unique role of VDAC2 in apoptosis (SFig. S5). Superposition of VDAC1 onto the VDAC2–BAX complex reveals several amino acid substitutions at the predicted interaction interface that alter the physicochemical properties of the binding surface and may contribute to the inability of VDAC1 to form a stable complex with BAX. The localization of this interface near residues previously implicated in both BAX and BAK regulation further supports the idea that VDAC2 has evolved specific structural features dedicated to the control of mitochondrial apoptosis.

Our findings provide a molecular explanation for the long-standing observation that VDAC2 is uniquely required for BAX-mediated apoptosis. We demonstrate that VDAC2 forms a stable complex with a membrane-inserted, activation-competent BAX conformation, thereby identifying a structural intermediate that links these observations. Recent studies have highlighted VDAC2 as an attractive therapeutic target for modulating apoptotic sensitivity in cancer and other diseases^15,48–50^. By defining the architecture of the VDAC2–BAX complex and the molecular determinants underlying its stabilization, our work provides molecular basis for the rational design of modulators targeting this interaction and for the development of new therapeutic strategies aimed at controlling mitochondrial apoptosis.

## Methods

### Production of recombinant proteins

Human VDAC2 was produced and purified as described in ^51^ with some modifications. Briefly, human 6His-VDAC2 gene was encoded in a pET22 plasmid containing an ampicillin resistance gene. The plasmid was cloned in *E. coli* BL21 cells. A single colony was inoculated in LB media and grown till OD 0.8 after which cells were induced with 1 mM IPTG and incubated at 37 °C for 4 hours. Cells were harvested and resuspended in lysis buffer (50 mM Tris pH 8.0, 2 mM EDTA, 20 % sucrose) followed by lysis by incubating for 10 minutes each in 1 mg/mL lysozyme, 0.6 % Triton X-100 and 10 µg/mL DNase. Cells were then broken by sonication at 50 % power with 50 % ON/OFF pulses for 10 minutes on ice. Lysates were centrifuged at 12,000 x g for 15 mins at 4 °C to pellet the inclusion bodies. The large, white inclusion bodies were resuspended and washed in wash buffer (20 mM Tris pH 8.0, 300 mM NaCl, 2 mM CaCl_2_) and re-centrifuged at 12,000 x g for 15 mins at 4 °C. The inclusion bodies were solubilized in equilibration buffer (20 mM Tris pH 8.0, 300 mM NaCl, 8 M urea,) by homogenizing and incubating for 1 hour at room temperature on agitation. Solubilized inclusion bodies were centrifuged at 150,000 x g in a Ti45 rotor for 1 hour at 4°C to remove all membrane contaminants. Supernatants were incubated with Ni-NTA resin in equilibration buffer. Contaminants were washed with equilibration buffer containing 20 mM imidazole and hVDAC2 was eluted with equilibration buffer containing 150 mM imidazole. Unfolded VDAC2 stocks in urea were concentrated to 250 µM in Amicon-10 centrifuge concentrators. Protein was incubated with 10 mM DTT for 10 minutes at room temperature before refolding.

Refolding was performed by slowly pipetting 1 mL of 250 µM unfolded, reduced hVDAC2 into 9 mL of pre-chilled refolding buffer (20 mM Tris pH 8.0, 150 mM NaCl, 1.5 % LDAO, 5 mM DTT) in a beaker with constant stirring at 4 °C. The mixture was incubated for 5-6 hours before transferring to a 3.5 kDa dialysis membrane and dialyzing against 1 L of dialysis buffer (20 mM Tris pH 8.0, 150 mM NaCl, 5 mM DTT) overnight at 4 °C. The next day, refolded protein was recovered and centrifuged at 150,000 x g in a Ti45 rotor for 60 minutes at 4°C to remove aggregates. The supernatant was concentrated in an Amicon-50 centrifuge concentrator to 0.5 mL and injected on a Superdex 200 increase 10/300 GL size exclusion column. The peak corresponding to monomeric, refolded VDAC2 was recovered, concentrated to 50 µM and stored at -80 °C in 20 % glycerol.

Human BAX gene cloned in pIVEX plasmid was obtained from Dr. Stephen Manon (IBGC, Bordeaux). BAX and BAX mutants were synthesized using a cell-free system of *E. coli* as described previously ^25^.

His-tagged MSP1D1 was produced and purified as previously described by ^52^. For His-tag cleavage, elutions of MSP1D1 were pooled and mixed with TEV protease at a 100:1 (wt/wt) ratio along with 1 mM DTT for 1 hour at room temperature. The mix was transferred to 3.5 MWCO dialysis tubing and dialysed against 100x volume of dialysis buffer (20 mM Tris pH 8.0, 300 mM NaCl, 1mM DTT, 10% glycerol) for 1 hour at room temperature followed by overnight at 4 °C. Cleaved MSP1D1 was re-purified on Ni-NTA resin, and the flowthrough and washes were collected, concentrated to 250 µM and stored at -80 °C. Cleavage efficiency was tested by SDS-PAGE and anti-His western blotting.

### Preparation of lipids and reconstitution of VDAC2 into nanodiscs

Chloroform solubilized chicken-egg phosphatidylcholine was dried under nitrogen gas. The lipid film was resolubilized in 20 mM Tris buffer pH 8.0 containing 50 mM sodium cholate at a concentration of 25 mM total lipid.

LDAO-refolded VDAC2 stocks at 50 µM were mixed with MSP1D1 and eggPC at a molar ratio of 1:12:780 giving an MSP:lipid ratio of 1:65 and a ratio of 6 nanodiscs/VDAC2. Required concentrations of proteins were mixed in a tube, along with buffer to maintain a final salt concentration < 200mM and glycerol concentration < 4%. 5 mM DTT was added during the reconstitution. The mixture was incubated at 4 °C for 1 hour before the addition of 100x detergent excess (wt/wt) of BioBeads SM-2 (in batches, every hour). The mixture was incubated overnight at 4 °C to remove all the detergent. The next day, nanodiscs were filtered via a 0.2 µ syringe filter and incubated on Ni-NTA resin on batch in 20 mM Tris buffer pH 7.2 containing 150 mM NaCl for 4 hours at 4°C on agitation. Empty nanodiscs were washed away with buffer containing 20 mM imidazole and VDAC2-containing nanodiscs were eluted with buffer containing 300 mM imidazole. Fractions were monitored at A_280,_ and protein-containing fractions were pooled and concentrated in an Amicon-50 concentrator to 0.5 mL. Concentrated VDAC2- containing nanodiscs were centrifuged at 20,000 x g for 30 mins at 4°C and loaded on a Superdex 200 increase 10/300 GL size exclusion column. VDAC2-nanodiscs eluting around 12 mL were collected and concentrated to 150 µM. Total protein was quantified using BCA assay. Quality of reconstitutions were verified by SDS-PAGE, dynamic light scattering and negative stain electron microscopy.

### Production and purification of VDAC2-BAX complexes

6His-VDAC2-containing nanodiscs were added to the cell-free reaction mixture at a final concentration of 3.5 µM, along with the plasmid encoding tag-free WT-BAX at a concentration of 15 µg/mL. The reaction mixture was dialyzed overnight at 28 °C with shaking at 120 rpm against a sufficient volume of filling mix, as described in ^25,36^.

The following day, the mixture was centrifuged at 20,000 × g for 20 min at 4 °C, and the supernatant was incubated with Ni-NTA resin in 20 mM Tris buffer (pH 7.2) containing 150 mM NaCl for 4 h at 4 °C. The resin was washed with the same buffer supplemented with 20 mM imidazole. For alkali extractions, the resin was incubated with 100 mM sodium carbonate (pH 10.0) for 30 min at room temperature. Excess carbonate was removed by washing as above. Finally, the complexes were eluted using buffer containing 300 mM imidazole. Elution fractions were monitored at 280 nm, and protein-containing fractions were concentrated to 0.5 mL using an Amicon-50 concentrator by centrifugation at 1,000 × g at 4 °C with intermittent pipetting.

The complexes were then centrifuged at 100,000 × g for 30 minutes at 4 °C prior to injection onto a Superdex 200 Increase 10/300 GL size-exclusion column. VDAC2–BAX complexes eluted at 11.3 mL. The presence of both proteins was confirmed by SDS–PAGE – western blotting.

### Co-immunoprecipitations

100 µL cell-free synthesis reactions containing either hVDAC1 or hVDAC2 nanodiscs with WT-BAX DNA were set up as described above. After overnight incubation at 28 °C with shaking at 120 rpm, the reactions were centrifuged at 20,000 × g for 20 min at 4 °C to remove aggregates.

The supernatants were diluted with binding buffer (20 mM Tris, pH 7.4, 250 mM NaCl), and 100 µL of Ni-NTA resin equilibrated in the same buffer was added to each sample. The resin was washed extensively with wash buffer (20 mM Tris, pH 7.4, 250 mM NaCl, 30 mM imidazole) after 2 hours of incubation at 4 °C. Purified proteins were eluted using the same buffer containing 300 mM imidazole. Eluted proteins were dialyzed against 2 L of dialysis buffer (20 mM Tris, pH 7.4, 150 mM NaCl) for 2 h at 4 °C. The samples were then recovered, supplemented with 5% glycerol, and split into four equal fractions. Each fraction was incubated overnight at 4 °C with agitation with the respective antibodies: anti-BAX 2D2 (Santa Cruz #20067, RRID:AB_626726), anti-BAX 6A7 (Santa Cruz #23959, RRID:AB_626728), anti-His (Proteintech #66005, RRID:AB_2857904), anti-VDAC1 (Proteintech #66345, RRID:AB_3672910) or a control (no antibody). The following day, 200 µL of protein G agarose resin was equilibrated with CoIP buffer (20 mM Tris, pH 7.4, 150 mM NaCl, 1 mM EDTA, 5% glycerol, 0.5 % NP-40), and 50 µL was added to each protein–antibody complex, followed by incubation for 4 h at 4 °C.

Samples were transferred to centrifuge spin columns and washed four times with 0.5 mL of CoIP buffer, followed by eight washes with 0.1x CoIP buffer (containing 0.05% NP-40). Finally, the resin was incubated with 30 µL of bromophenol blue sample buffer containing SDS and β-mercaptoethanol for 30 min at room temperature without heating. Eluates were collected by centrifugation and 15 µL of sample each was loaded and run on 15% SDS-PAGE gels. Gels were transferred to nitrocellulose membranes at 80 mV for 50 minutes at 4 °C. Membranes were blocked with 5% milk in 1x PBS-T for BAX antibodies and 3% BSA in 1x PBS-T for VDAC antibodies. BAX antibody (Abcam #182734, RRID:AB_3099426, diluted 1:10,000 in PBS), VDAC2 antibody (Abcam #316107, diluted 1:1000 in PBS), and VDAC1 antibody (CST #4661, RRID:AB_10557420, diluted 1:1000 in PBS) were used for revelation.

### Cryo-Electron Microscopy

#### Cryo-EM of VDAC2–BAX nanodiscs

Purified VDAC2–BAX nanodiscs were concentrated to 0.4 mg/mL in 20 mM HEPES, 150 mM NaCl (pH 7.4). Samples were applied to graphene oxide-coated Quantifoil R1.2/1.3 300-mesh copper grids prepared as previously described52. Prior to sample application, grids were plasma-cleaned at 50 mA for 1 min. A 4.5 μL aliquot of sample was applied to each grid using a Vitrobot Mark IV (Thermo Fisher Scientific). Grids were blotted for 3 s with a blot force of 2 under 100% relative humidity and plunge-frozen in liquid ethane. Cryo-EM data were collected on a Glacios transmission electron microscope (Thermo Fisher Scientific) operated at 200 kV and equipped with a Falcon 4i direct electron detector and a Selectris X energy filter. Movies were recorded in electron-counting mode at a nominal magnification of 120,000×, corresponding to a calibrated pixel size of 0.90 Å/pixel. A total exposure of 40 e−/Å² was fractionated into 40 frames and collected over a defocus range of −0.2 to −2.0 μm using a 10 eV energy-filter slit.

#### Cryo-EM of VDAC2 nanodiscs

Purified VDAC2 nanodiscs were concentrated to 0.6 mg/mL in 20 mM HEPES, 150 mM NaCl (pH 7.4). Samples were applied to glow-discharged Cu/Rh Quantifoil R1.2/1.3 grids using a Vitrobot Mark IV (Thermo Fisher Scientific). A 3 μL aliquot of sample was applied to each grid, blotted for 3 s with a blot force of 2 under 100% relative humidity, and plunge-frozen in liquid ethane. Cryo-EM data were collected at the European Synchrotron Radiation Facility using a Titan Krios G4 transmission electron microscope (Thermo Fisher Scientific) operated at 300 kV and equipped with a cold field emission gun (CFEG), fringe-free illumination and an in-column Gatan Quantum LS energy filter with a 20 eV slit width. Movies were acquired at a nominal magnification of 130,000×, corresponding to a calibrated pixel size of 0.66 Å/pixel. Data were collected without a phase plate and recorded with an exposure time of 1.08 s, fractionated into 40 frames, at a dose rate of 16.2 electrons/pixel/s, corresponding to a total exposure of 40.5 e−/Å² per movie. Images were acquired over a defocus range of −1.0 to −2.2 μm in 0.2 μm increments. The objective lens had a spherical aberration coefficient (Cs) of 2.7 mm.

### Image processing

#### The VDAC2–BAX dataset

Image processing was performed using Scipion (RRID:SCR_016738). A total of 5,502 movies were collected. Beam-induced motion correction and frame alignment were carried out using MotionCor2 (RRID:SCR_016499), while CTF parameters were estimated with CTFFIND. A total of 1,989,848 particles were automatically picked using crYOLO. Three consecutive rounds of 2D classification were then performed to remove false positives and low-quality particles. After particle cleaning, 31,927 particles were retained for ab initio reconstruction in cryoSPARC (RRID:SCR_016501) to generate initial 3D models. The resulting initial volumes were further refined through several rounds of Heterogeneous Refinement in cryoSPARC, allowing the identification of the class corresponding to particles containing two VDAC pores and an extra density. A total of 12,498 particles belonging to this class were subsequently subjected to Non-uniform Refinement followed by Local Refinement in cryoSPARC, yielding a final reconstruction with an overall resolution of 8.1 Å, based on the gold-standard Fourier Shell Correlation (FSC) criterion. The final density map was sharpened using DeepEMhancer.

#### The VDAC2 dataset

Image processing was performed using CryoSPARC (RRID:SCR_016501). A total of 3,974 dose-fractionated movies were subjected to Patch Motion Correction followed by Patch CTF Estimation using the default parameters. Initial particle picking was performed using the Blob Picker with a minimum particle diameter of 130 Å and a maximum particle diameter of 300 Å. Particles were extracted using a box size of 300 pixels and subjected to three rounds of reference-free two- dimensional (2D) classification to remove false positives and poorly defined particles. The resulting high-quality classes were used for template-based particle picking, generating a dataset of approximately 56,000 particles. An initial three-dimensional model was generated by single-class Ab Initio Reconstruction and iteratively improved through successive rounds of Ab Initio Reconstruction and Heterogeneous Refinement to remove remaining contaminants and improve particle homogeneity. The final particle stack was refined using Non-Uniform (NU) Refinement to yield a three-dimensional reconstruction with an overall resolution of 6.6 Å, as estimated by the gold-standard Fourier shell correlation (FSC) criterion at the 0.143 threshold.

### Cysteine crosslinking VDAC2-BAX complexes

Single cysteine mutants of VDAC2 and BAX were produced in a 100 µL cell-free system as described above. The mixtures were centrifuged at 20,000 x g for 20 minutes at 4°C and purified on Ni-NTA resin as explained above. Complexes were eluted in 0.5 mL of buffer containing 300 mM imidazole. Complexes were subsequently split into two fractions and treated either with 1 mM of copper phenanthroline or 2.5 mM of TCEP for 30 mins at 30 °C with agitation at 650 rpm. Reactions were quenched by adding 10 mM EDTA for 10 minutes at 30 °C at 650 rpm. Finally, samples were incubated with 2.5 mM NEM for 10 minutes at 30 °C to quench any free cysteines. Proteins were precipitated using methanol/chloroform and loaded on 15 % SDS-PAGE gels for western blotting against BAX and VDAC2 as explained above, using BAX antibody (Santa Cruz #20067, diluted 1:3000 in PBS) and VDAC2 antibody (Abcam #316107, diluted 1:1000 in PBS).

### NEM-PEG accessibility labelling

Single cysteine mutants of BAX were produced in a 100 µL cell-free system as described above either in the presence of wt-6His-VDAC2 nanodiscs or alone. Samples of BAX alone were not purified on resin, while VDAC2-containing samples were purified on Ni-NTA resin as explained above, this time however, recovering the flowthrough of the Ni-NTA resin. Complexes were eluted in the same volume of buffer as the recovered flowthrough. Samples of BAX alone, VDAC2-BAX flowthrough and elution were labelled with 0.2 mM mPEG-MAL MW 5k (Creative PEGWorks #PLS-234) for 1 min on ice followed by quenching with 20 mM NEM for 10 mins at RT on agitation. Equal volumes (5-10 % of total CFS) of NR and EL was loaded and run on 15% SDS-PAGE gels followed by revelation with BAX antibody (Santa Cruz #20067 2D2, diluted 1:3000 in PBS).

### VDAC channel reconstitution in planar lipid membrane and electrical measurements

VDAC channel measurements were carried out by reconstituting LDAO-folded wt-VDAC2 into a planar lipid membrane built using the modified Montal-Mueller technique. Membranes were prepared from neutral diphytanoyl phosphatidylcholine (DPhPC) or soybean polar lipid extract (PLE). Membranes were purchased from Avanti Polar Lipids. Aliquots of 10–20 µL of 5 mg/mL total lipids were added on top of each salt solution in two 1.8 mL compartments (so-called *cis* and *trans*) of a Teflon chamber. The two compartments were separated by a 15 µm-thick Teflon film with a 70–100 µm diameter orifice where the membrane is formed. The orifice was pre-treated with a 3% solution of hexadecane in pentane. After pentane evaporation, the level of solutions in each compartment was raised above the hole using syringes, so the planar bilayer could form by apposition of the two monolayers. The chambers were filled with 5 mM HEPES buffer pH 7.4 containing either 0.15 M or 1 M KCl and 5 mM DTT.

VDAC channel insertion was achieved by adding 0.5 µL of LDAO-refolded VDAC diluted at 0.003 mg/mL in buffer containing 10 mM Tris, 50 mM KCl, 1mM EDTA, 15% (v/v) DMSO, 2.5% (v/v) Triton X-100, pH 7.0 into the *cis* compartment. Upon reconstitution of a single VDAC2 channel, currents were recorded for some time to ensure that the channel was stable and no more channels got inserted. As previously reported for VDAC reconstituted in planar lipid bilayers, most channels insert with a uniform orientation, with the *cis* side of the chamber presumably corresponding to the cytosolic face of VDAC^53,54^. After VDAC insertion, BAX was added to the *cis* compartment in the bulk. After addition of BAX, a triangular wave with an amplitude of 5 mV Vpp (peak-to-peak voltage) and a frequency of 5 mHz was applied for 30-60 minutes (SFig. S2A). The triangular wave is considered to assist in the interaction of BAX with the VDAC channel. To assure reproducibility, the experiments were repeated a minimum of 3 times. Experiments were also performed in different lipids with different salt concentrations.

The membrane potential was applied using Ag/AgCl electrodes in 2 M KCl/1.5% agarose bridges assembled within standard 200-μl pipette tips. Potential is defined as positive when it is greater at the side of protein addition (*cis* side). Current recordings were performed using an Axopatch 200B amplifier (Axon Instruments, Inc.) in voltage-clamp mode. Data from the amplifier were filtered by an integrated low pass Bessel filter at 10 kHz, digitized with a Digidata 1440A (Molecular Devices, Sunnyvale, CA) at a sampling frequency of 50 kHz and analyzed using pClamp 10.7 software (Molecular Devices, Sunnyvale, CA, RRID:SCR_011323). The chamber and the head stage were isolated from external noise sources with a double metal screen (Amuneal Manufacturing Corp., Philadelphia, PA). The set-up described can measure currents of the order of picoamperes or above with a time resolution below the millisecond.

### Alphafold predictions and molecular dynamics simulations

AF3.0 predictions were performed on the Alphafold web server using VDAC2 (UniProt : P45880), BAX (UniProt : Q07812) and 50 molecules of palmitic acid ligands. No specific MSAs or templates were provided. Models were performed in triplicates yielding a total of 15 models.

The first-ranked AlphaFold predicted structure was selected for coarse-grained MD simulations using the MARTINI 3.0 force field (v3.0.b.3.2). An elastic network was applied to maintain contacts between residues separated by less than 9.0 Å, using a force constant of 500 kJ·mol⁻¹·nm⁻². The coarse-grained system, including 200 lipids in a 90:10 POPC:POPG mixture, was built using the CHARMM-GUI interface.

The system was subjected to two consecutive steepest-descent energy minimizations (5000 steps each), followed by five equilibration runs. The first minimization employed a soft-core potential applied to both Coulombic and van der Waals interactions in order to soften interactions between overlapping particles.

The equilibration procedure consisted of successive 10 ps stages during which the integration timestep was gradually increased from 2 to 20 fs. The system was directly heated to 303.15 K using separate thermal baths for protein, membrane, and solvent groups, coupled with the v-rescale thermostat using a coupling constant (τt) of 1 ps. Pressure was maintained at 1.0 bar using the Berendsen semi-isotropic barostat with a coupling constant (τp) of 5 ps.

During equilibration, position restraints were applied using a progressively decreasing force constant scheme ranging from 1000 to 50 kJ·mol⁻¹·nm⁻² for protein backbone beads and from 200 to 10 kJ·mol⁻¹·nm⁻² for lipid phosphate groups.

Production simulations were subsequently carried out for 10 µs without positional restraints. Pressure coupling was then switched to the Parrinello–Rahman semi-isotropic barostat with τp =12.0 ps and a compressibility of 3×10−4 bar⁻¹ in both membrane dimensions. The simulations were run in triplicates, using different starting lipid configurations.

### Pore conductivity

To compute the conductivity of VDAC during the simulations, we first selected 1,000 configurations from each simulation. These configurations were aligned along the Z-axis, centered, and fitted to the VDAC protein backbone beads. The structures were then transformed back to an all-atom representation using the backward procedure ^55^.

We applied the Hole program ^56^ to each configuration, initiating the analysis from nine distinct ion positions within the channel center to sample a broad range of possible ion pathways. Conductivity was determined based on the atomic radii defined in the CHARMM force field. For each attempt, the narrowest part of the pathway was identified, and the conductivity was estimated from the averaged value of the nine determinations.

This procedure was repeated for both the VDAC channel alone and the VDAC-BAX complexes, yielding a time-dependent measurement of conductivity over the MD timescale. Notably, this approach estimates an average conductivity of 4.6 nS/M of KCl for the fully open channel, which aligns closely with experimental measurements of 4 to 4.5 nS reported across various organisms ^57^. To compare the conductance ability of VDAC in the presence or absence of BAX, we computed an open probability based on a threshold conductance of 1 nS/M of KCl.

### Cross-linking Mass Spectrometry (XL-MS)

WT-VDAC2 and WT-BAX complex produced in a 2 mL cell-free system as described above was subjected to Ni-NTA purification and gel filtration chromatography in 20 mM HEPES buffer (pH 8.0), 150 mM NaCl as explained above. The shouldered peak of the chromatogram at 11.3 mL was recovered, concentrated to 200 µL and quantified using BCA assay, Roughly 200 µg total protein was subjected to lysine crosslinking using isotopically labeled Di-SuccinimidylSuberate d0/d12 (DSS-H12/D12, Creativemolecules Inc.) at a molar ratio of 1:1 crosslinker to lysines for 30 min at 30 °C at pH 8.0 while shaking at 650 rpm in a Thermomixer (Eppendorf), as described previously ^58^. The reaction was quenched with Tris buffer at a final concentration of 50 mM for 15 min at 30 °C at 650 rpm. The cross- linked sample was precipitated in methanol/chloroform to remove lipids and resuspended in 200 μL of 8 M Urea. The samples were then reduced, alkylated, and digested with trypsin (Promega), at 1:30 (E:S ratio). Digested peptides were purified by solid-phase extraction (50-mg SepPak cartridges, Waters), and cross-linked peptides were enriched by size exclusion chromatography using an ÄKTAmicro chromatography system (GE Healthcare) equipped with a Superdex^TM^ Peptide 3.2/30 column. Absorption levels at 215 nm of collected fractions were used to normalize peptide concentration prior to analysis by liquid chromatography-tandem mass spectrometry (LC-MS/MS).

LC-MS/MS analysis was performed on an Orbitrap Eclipse Tribrid mass spectrometer. Peptides were separated on a Vanquish Neo system (Thermo Scientific) at a flow rate of 300 nL/min over a 60 min gradient optimized for each fraction (3 % to 5-12 % for 4 min, 5-12 % to 20-32 % in 35 min, 20-32 % to 28-36 % in 10 min, 28-36 % to 80 % acetonitrile in 0.1 % formic acid in 1 min, 80% acetonitrile in 0.1 % formic acid for 10 min). Full scan mass spectra were acquired in the Orbitrap at a resolution of 120,000, a scan range of 400–1650 *m/z*, and a maximum injection time of 100 ms. Most intense precursor ions (intensity ≥5.0 × 10^3^) with charge states 2–8 and monoisotopic peak determination set to “peptide” were selected for MS/MS fragmentation by CID at 35 % collision energy in a data-dependent mode. The duration for dynamic exclusion was set to 45 s. MS/MS spectra were analyzed in the Ion Trap at a rapid scan rate and maximum injection time in dynamic mode.

The experiment was performed in biological duplicates which were additionally measured as technical duplicates each. Crosslinks were identified and filtered stringently using xProphet to an FDR ≤ 0.05 and the data was visualized using xiNET.

## Acknowledgements

This work was supported by the ANR Apo_Mito project (L.B.), the Aix-Marseille Université AMX-21- PEP-022 grant (L.B.), and a PhD fellowship from Aix-Marseille Université (V.R.). This work benefited from access to the Instruct Image Processing Center, an Instruct-ERIC Centre, with financial support provided under PID 41299. Computational resources were provided by GENCI–IDRIS under Grants 2021-A0110707044, 2022-A0110707044 and 2023-AD010707044R1. F.S. is grateful for funding by the German Research Foundation (project numbers 496470458, 516836828, 538898546 and TRR353/1 – 471011418).

## DATA AVAILABILITY

The mass spectrometry data (raw files and xQuest, search output files) have been deposited to the ProteomeXchangeConsortium via the PRIDE^59^ repository under the accession number PXD080454. (Reviewer access details Project accession: PXD080454 and Token: FlAwoW1I1Zf0). Molecular dynamics trajectories, input files, and the structural models generated in this study will be deposited in Zenodo (https://doi.org/10.5281/zenodo.21157499). Source data underlying all figures, including uncropped gels and western blots, will be provided with the manuscript.

## Notes

### Competing Interest Statement

The authors have declared no competing interest.

